# Occurrence of a ‘*Candidatus* Phytoplasma pruni’-Related Strain Associated with Commercial Poinsettias in Turkey

**DOI:** 10.1101/2024.11.12.623157

**Authors:** Kemal Benlioglu, Umit Ozyilmaz, Huseyin Faruk Yurttas

## Abstract

The poinsettia (*Euphorbia pulcherrima* Willd. ex Klotzsch.) is one of Turkey’s most important potted ornamental crops. Potted poinsettia plants exhibiting smaller-sized brilliant red bracts against the green leaves with free-branching were collected from commercial florists in three provinces of Turkey. In iPhyClassifier analysis, the query 16SrDNA sequences of our isolates shared 99.58% identity with that of the *Candidatus* Phytoplasma pruni. Our leaf samples consistently yielded a product of 0.7 kb by PCR with primers (KFD4F/KFD4R), which was newly designed based on X-Disease group (16SrIII) of phytoplasma secY sequences available in GenBank. The partial 16S rRNA sequences were used for phylogenetic analyses covering 41 taxa from the group of 16SrIII^’^ Candidatus Phytoplasma’ strains, including 14 strains causing disease on poinsettia worldwide. The sequences that caused the same disease in poinsettia plants in Finland, the USA, Mexico, Taiwan, Japan, Korea, and Canada, including three strains from Turkey, formed a mono-phylogenetic group with a support value of 87%.

## Introduction

The poinsettia a member of the large and diverse family Euphorbiaceae, originated in southern Mexico and northern Guatemala. In its native habitat, this species is a winter-flowering shrub that grows over 3 m and is a common landscape plant. The sap is milky and may produce dermatitis in susceptible individuals. It was cultivated by the Aztecs of Mexico, near present-day Taxco. The brilliant ‘flower’ was beloved by natives and their kings as a symbol of purity. Poinsettias were first introduced into the United States in 1825 by Joel Robert Poinsette while he served as the first U. S. Ambassador to Mexico. The poinsettia was widely distributed across Europe by the mid-19th century. It enjoyed great popularity because it succeeded in North America (Taylor et al., 2011). The poinsettia is one of Turkey’s most important potted ornamental crops and has enjoyed great popularity since being called Atatürk’s Flower in 1929 (Durak, 2014).

In 1995, Lee et al. reported that these poinsettias’ commercially popular compact appearance and demanded economic importance were due to a free branch-inducing factor, the PoiBI phytoplasma. The phytoplasma was characterized using polymerase chain reaction-restriction fragment-length polymorphism (PCR-RFLP) analysis, and the predominant strain present was shown to belong to the 16S rDNA group III, which includes X-disease and related phytoplasmas (Lee et al., 1995; 1997).

To investigate the genetic identity and phylogenetic relationships of phytoplasma strains infecting poinsettia plants in Turkey, focusing on identifying their association with the 16SrIII X-Disease group. The study aims to characterize the isolates collected from three provinces using 16S rRNA sequencing and secY-specific PCR to enhance understanding of the pathogenic phytoplasma strains affecting poinsettia and contribute to global data on the spread and evolution of these pathogens.

## Material and Methods

In September 2017, potted poinsettia plants exhibiting showy, scarlet bracts (leaflike structures attached just below flowers) surrounding a central cluster of tiny greenish-yellow flower clusters were collected from commercial florists in three provinces (Ankara, Antalya, and Aydın) of Turkey (Fig. 1). Potted poinsettias were dwarf and lovely compact-sized plants with free-branching and a multi-flowered canopy, exhibiting smaller size brilliant red bracts against the green leaves. The three sampled poinsettias with commercial popularity had the same appearance as previously described phytoplasma-infected plants (PoiBI) with a free-branching morphotype characterized by weak apical dominance and many axillary shoots ‘flowers’ by Lee et al. (1997). DNA was extracted from flowering-stage poinsettia plants using leaf midribs and petioles of about 5 or 6 leaves after bracts from the top to the downward of each plant with a modification of the DNA isolation procedure of Ahrens and Seemüller (1992). For extraction, approximately 0.5 g of fresh leaf venal tissue was cut into small pieces and was grounded after incubation for 10 min in 5 ml of ice-cold grinding buffer (125 mM potassium phosphate, 30 mM ascorbic acid, 10% sucrose, 0.15% bovine serum albumin (BSA), 2% polyvinylpyrrolidone (PVP-40, pH 7.6) in a prechilled mortar. The extract was passed through two layers of sterile cheesecloth, followed by filtration through Whatman 1 and 5 filter papers. The clear extract was then filtered through a 0.45 μm Millipore filter (Millipore, USA) due to the pleomorphic morphology of phytoplasma enabling it to pass through the 0.45 μm pores. The homogenate was centrifuged at 4° C for 25 min at 14600 g. The pellet was resuspended in 1.5 ml of warm (60° C) extraction buffer according to Doyle (1991) containing (2% CTAB, 1.4 M NaCI, 0.2% 2-mercaptoethanol, 20 mM EDTA, 100 mM Tris-HCI, pH 8.0) and was incubated at 65° C for 30 min. The lysate was extracted with equal chloroform/isoamyl alcohol (24:1, v/v). After centrifugation, the aqueous layer was precipitated with a two-thirds volume of -20° C isopropanol and centrifuged at 14600 g with a microfuge. The pellet was washed with 70% ethanol, dried under vacuum, and dissolved in 100 µl TE buffer (10 mM Tris, one mM EDTA, pH 8.0).

**Fig. 1.**
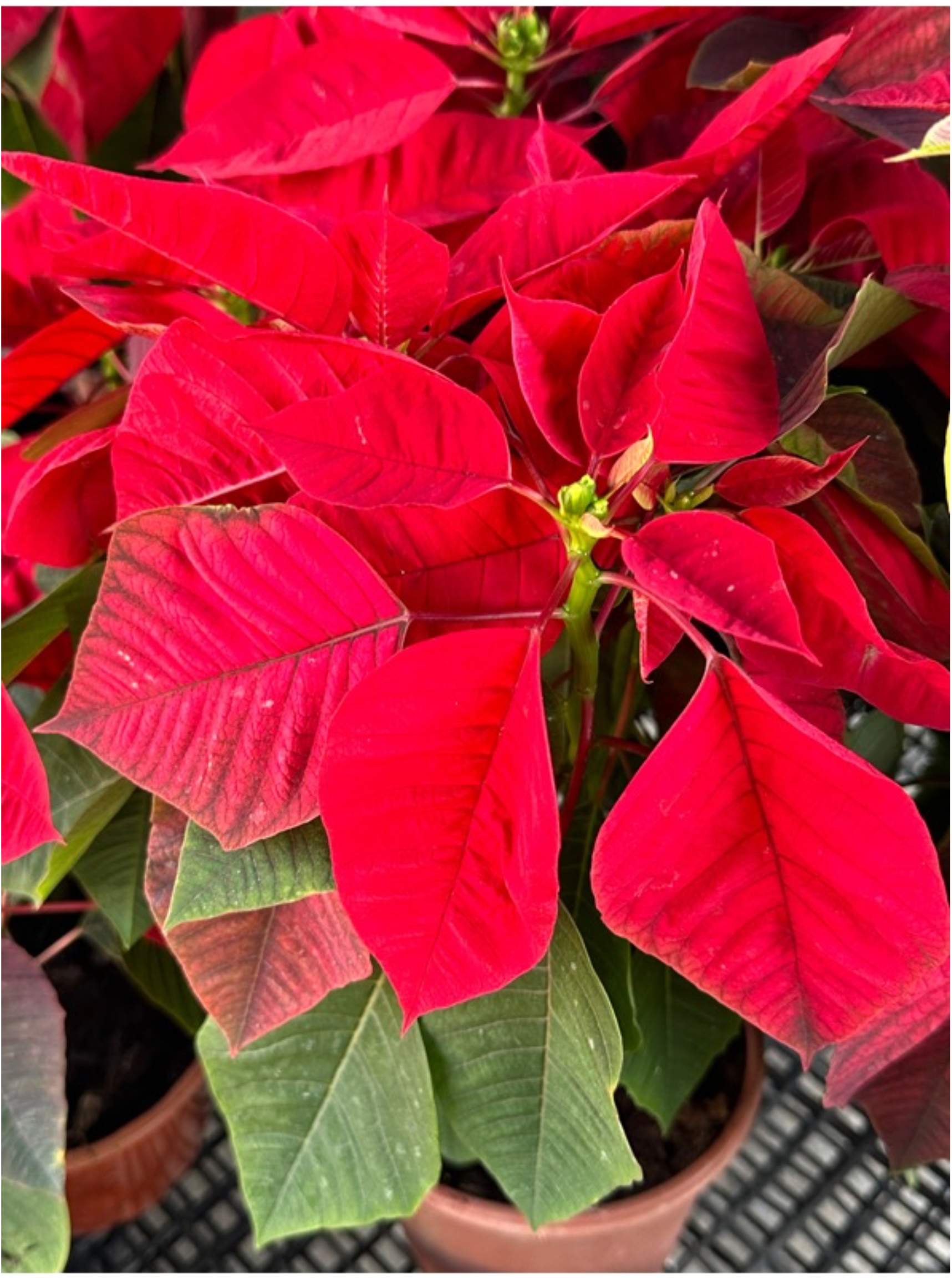
Poinsettia plant (*Euphorbia pulcherrima*) exhibiting showy, scarlet bracts smaller lovely compact-sized plants with free-branching canopy against the green leaves.

For phytoplasma detection, the universal primer set P1 / P7 (Schneider et al., 1995) was used for the first (direct) amplification, and R16F2n/R16R2 was used for the second (nested) amplification (Gundersen & Lee, 1996), targeting the 16S rRNA gene sequence of 1.8 and 1.2 kb in length, respectively. Reactions were cycled in a C1000 thermal cycler (Bio-Rad Inc.) with DreamTaq DNA Polymerase and PCR master Mix (Thermo Scientific). Direct PCR products were diluted with sterile distilled water (1:20) before the nested PCR. Parameters for PCR conditions were as follows: initial denaturation at 95° C for 3 min, followed by 35 cycles of 94° C for 10 s, 55° C (60° C for nested PCR) for 15 s, and at 72° C for 10 s with the final extension set at 72° C for 5 min. After amplification, a 4-μl aliquot from each sample was electrophoresed in a 1% agarose gel and visualized by staining with ethidium bromide and UV illumination. The sequencing of PCR products was performed by Macrogen Service (Seoul, Korea) and submitted to the GenBank “NCBI” under the accession numbers MN398473, MN398474, and MN398475. Three molecular diagnostic tools, PCR-RFLP, PCR amplification of a specific target gene, and phylogenetic analysis, were used to Identify phytoplasma strains causing PoiBI. For PCR-RFLP, the resulting 16S rRNA gene sequences were subjected to virtual RFLP analysis via the iPhyClassifier software (Zhao et al., 2009). Secondly, the identification and confirmation of phytoplasma strains, the secY gene, encoding a protein translocase subunit, was amplified with KFD4F (5’-GAA AGA ATG GCA AGA ACA AGG -3’) and KFD4R (5’ CCA GGT TTT ACC CCT TCT AAG-3’) primer pair, which was newly designed based on X-Disease group (16SrIII) of phytoplasma secY sequences available in GenBank. PCR was performed with the secY-specific primer pairs using the same reagents and under the same conditions except at annealing temperatures of 53° C as previously described and visualized on an agarose gel. Phylogenetic analysis was done by aligning the partial 16S rDNA sequences from 3 strains (MN398473, MN398474, and MN398475) and the sequences obtained from the GenBank to compile datasets. Multiple sequence alignments were performed in MEGA X (Kumar et al., 2018). Phylogenetic analysis was performed using MEGA X software using the Clustal-W algorithm and neighbor-joining method with 1000 bootstrap replications.

## Results and Discussion

In iPhyClassifier analysis, the query 16S rDNA sequences of our isolates (AC1, AC2, and AC3) shared 99.58% identity with that of the “Candidatus Phytoplasma pruni rrnA’ reference strain (GenBank accession: JQ044393) and noticed that the phytoplasma under study was a “Candidatus Phytoplasma pruni’ rrnA’-related strain. This result is in agreement with those of Rosete et al. (2021) and Lee et al. (1997) in the US, where phytoplasmas associated with a free-branching morphotype characterized by weak apical dominance and many axillary shoots and ‘flowers’ were enclosed in the 16SrIII group (Lee et al., 1998; 2000; Pondrelli et al., 2002). Other research in Korea (Chung & Choi, 2010) revealed that symptomatic poinsettia had 99.6 identity with U.S. PoiBI isolate FJ376625, indicating poinsettia stem flat disease is caused by PoiBI. However, we could not observe any symptoms like flat stems and fascicles and an abnormal number of apexes resulting in a cockscomb stem form, leaf narrowing with curling of bracts as described in Korea.

All leaf samples from three symptomatic plants consistently yielded a product of 0.7 kb by PCR with our newly designed primer pairs. It is known that a single copy of the secY gene is present in all known phytoplasma genomes. It is one of the most variable among the phylogenetic markers used so far for differentiation of phytoplasma strains and was found to be more efficient than other gene markers for differentiation and classification of phytoplasmas (Lee et al., 2006; Lee et al., 2010). The designed Primer pairs (KFD4F/KFD4R) were blasted against the nucleotide collection BLAST database (nr), which is limited to sequences from bacteria. The in-silico primer validation by Primer-BLAST also indicated that these oligonucleotides could detect all Candidatus phytoplasma pruni and all 16SrIII group variants available in the GenBank dataset and those reported by Lee et al. (2010) and Davis et al. (2015).

The partial 16S rRNA sequences (approximately 1200 bp) were used for phylogenetic analyses from 41 taxa from the group of 16SrIII’ Candidatus Phytoplasma’ strains, including 14 strains causing disease on poinsettia worldwide and outgroup (Acholeplasma palmae) obtained from GenBank. The model for phylogenetic analysis was selected by fitting the Neighbor-Joining method and the Tamura-Nei model (Tamura & Nei, 1993), choosing the lowest BIC scores (Bayesian Information Criterion). The phylogenetic tree was constructed with 1000 bootstrap steps (Fig. 2). There were 1203 positions in the final dataset. Evolutionary analyses were conducted in MEGA X (Kumar et al., 2018; Stecher et al., 2020).

**Fig. 2.**
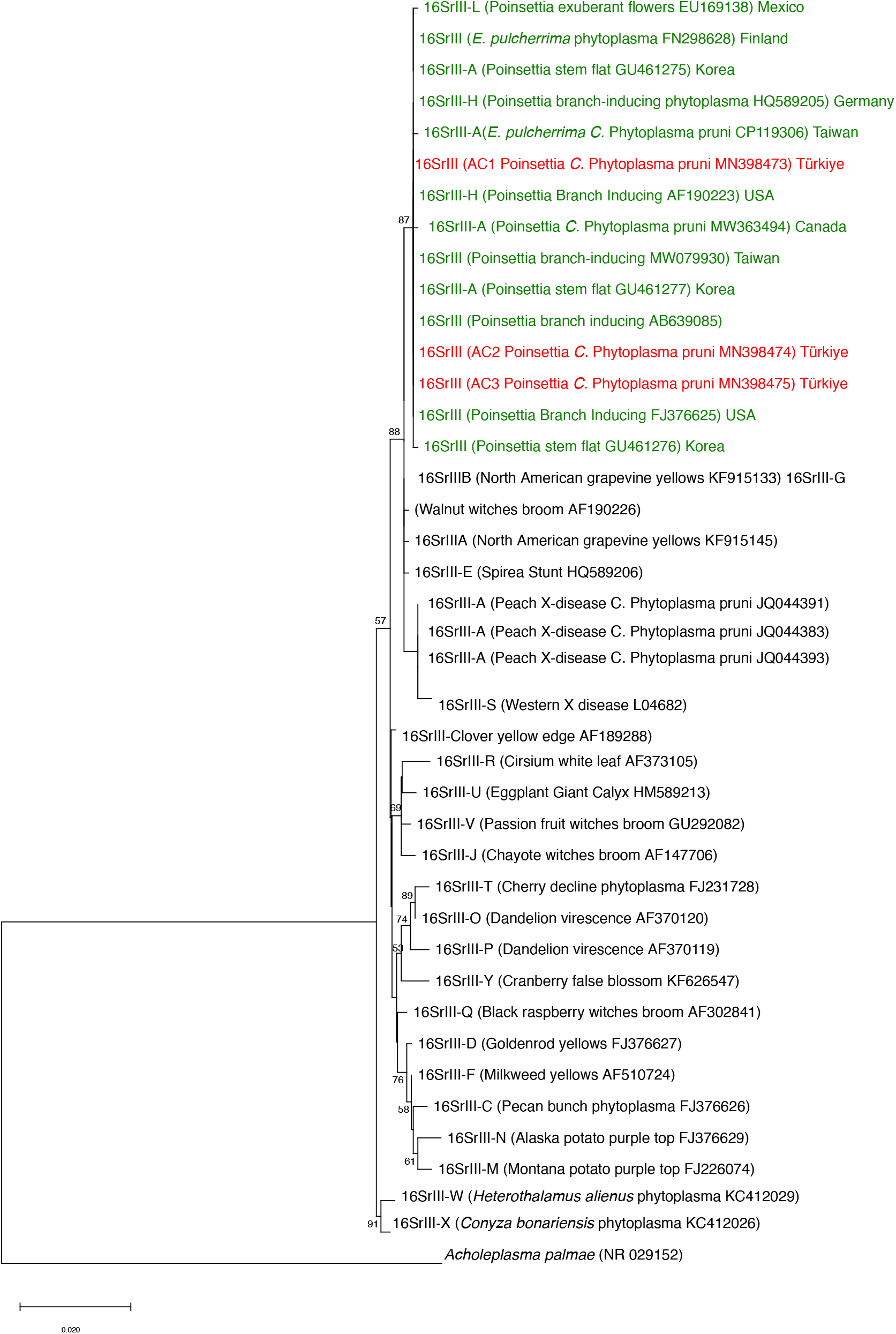
A neighbor-joining tree derived from a Clustal-W sequence alignment of 16SrRNA sequences of the associated 41 phytoplasma strains including *Acholeplasma palmae* as an out-group. The percentage of replicate values clustered in the associated taxa in the bootstrap test (1000 replication) is shown next to the branches, except for less than 50%. The evolutionary distances were computed using the Tamura-Nei method and are in the units of the number of base substitutions per site. Phylogenetic sequence data analyses were done using MEGA X (Kumar et al., 2018).

The first clade, with a support value of 87, contained partial 16SrRNA sequences from the Candidatus phytoplasma, which caused the same disease with different manifestations or similar manifestations but different names in poinsettia plants in Finland, USA, Mexico, Taiwan, Japan, Korea, and Canada, including three strains (AC1, AC2, AC3) from Turkey (Fig. 2). By investigating this rDNA from phytoplasmas from 20 poinsettia cultivars, Lee et al., (1997) found that the predominant type of phytoplasma was a member of the 16SrDNA group III and is most closely related to x-disease and spirea stunt. Phylogenetic tree based on the 16S rRNA gene sequence of the Ontario poinsettia phytoplasma (MW363494) in Canada indicated that all 16SrIII group phytoplasma formed a monophyletic clade with strong bootstrap support of 100%, and three poinsettia phytoplasma strains clustered a subclade with 66% support within the main clade (Rosete et al., 2021). However, further studies are needed on the subgrouping system using the 16Sr RNA gene for adequate branch support and significant geographic and strain relevance. This is the first report of the occurrence of a 16SrIII phytoplasma “Candidatus Phytoplasma pruni” strain in E. pulcherrima in Turkey in commercial poinsettias. Poinsettias with red and green foliage are the most popular and high-value potted plants worldwide, with more than 2 million sold yearly in Turkey. Free-branching cultivars have been in Turkey for a long time, but no western X-like disease has been reported in the country. It is unknown whether a vector for PoiBI is present in Turkey. However, it is worth noting that a free-branching agent is graft-transmissible (Dole & Wilkins, 1992). Preil and Ebbinghaus (1998) also demonstrated the transmission to Euphorbia fulgens (scarlet plume) by grafting with poinsettia as the rootstock, while the Branch-inducing phytoplasmas from poinsettia were also transmitted by dodder to the crown of thorns (Euphorbia milii) (Nicolaisen, 2004). Given the increasing awareness of phytoplasmas as an emerging pathogen, handling poinsettias with utmost care is crucial to prevent the potential phytosanitary risk of phytoplasma spread within the country. Our findings underscore the need for careful action and heightened vigilance in plant pathology and agriculture regarding phytoplasma diseases.

## Acknowledgments

This paper was supported by a project, “ADÜ-BAP ZRF-18030” from the Aydin Adnan Menderes University, Turkey.

